# Novel and simple simulation method to design and development of antisense template

**DOI:** 10.1101/2022.09.30.510283

**Authors:** Devendra Vilas Deo, Nawaj Shaikh

## Abstract

Antisense technology is emerging as potential therapeutics against lethal infections. Basically, Antisense-mRNA complex inhibits the protein translation of pathogens and thus it is used for treatment. Based on previous online tools and literatures and difficulties for designing antisense template, finding high conserved regions from large number of long sequences, by taking all those factors in consideration, we proposed new innovative offline target simulation methods i.e. Deletion of unwanted region from viral sequence alignment (DURVA) and Most frequent region (MFR) for designing and developing antisense template from large number of long sequence or genomic data. Based on current pandemic crisis and long genomic sequence of SARS-CoV-2, we chose coronavirus for simulation. Initially, we hypothesized that DURVA-MFR would find stable region from large annotated sequencing data. As per Chan et.al. guidelines for antisense designing and development, we designed couple of algorithms and python scripts to process the data of approximately 30kbp sequence length and 1Gb file size in short turnaround time. The steps involved were as: 1) Simplifying whole genome sequence in single line; 2) Deletion of unwanted region from Virus sequence alignment(DURVA); 3)Most frequent antisense target region(MFR) and 4)Designing and development of antisense template. This simulation method is identifying most frequent regions between 20-30bp long, GC count≥10. Our study concluded that targets were highly identical with large population and similar with high number of remaining sequences. In addition, designed antisense sequences were stable and each sequence is having tighter binding with targets. After studying each parameter, here we suggested that our proposed method would be helpful for finding best antisense against all present and upcoming lethal infection.The initial design of this logic was published in Indian Patent Office Journal No.08/2021withApplication number202121005964A.

**Simple summary:** The antisense development is state of the art for modern therapeutics. There are number of online soft-wares and open sources for designing of antisense template. But all other tools did not consider frequency as major factor for designing antisense. Also; all sources excepting our simulation approach does not process large file or long sequences. Therefore; we designed an offline innovative simulation method which deletes the unwanted region from sequences and stores the data which are fulfilled antisense criteria. Further; the calculation of frequency from these short listed target regions; the most frequent region is desire antisense target and further antisense template will be designed according to Watson-Crick model. This article explained all information about how our new approach is best for designing antisense template against SARS-CoV-2 and many lethal infectious viruses etc.

## Introduction

Modern medicine looking towards antisense technology. Antisense is short template around 20-30bp and it binds complimentary to host mRNA. Antisense-mRNA complex inhibits the protein translation. Thus; antisense involves in regulation of gene expressions. This is a modern approach to inhibit the viral protein translation in case of viral infection(1–3). We enrolled new and popular pandemic virus nowdays for study purpose and to evaluate the efficiency and accuracy of the our self-derived methodology for long and complex genomic sequences which was not easy to defined conserved and stable targets because of long and complex genomic sequence structure. The Corona virus belongs to family *Coronaviridae* and genera Betacoronavirus. This is positive single stranded genomic RNA virus about 29Kbp long and act as mRNA within host cell. Generally, the strains of virus attacks on respiratory and other systems and it binds to Angiotensin-converting enzyme 2 (ACE-II enzyme) receptors present on respiratory system cells and multiply within the host cells.This persistent infection may cause lungs damage.(4) This was another reason to choose SARS-CoV-2 in addition with assess the accuracy and effectiveness of our self-designed method.

In case of COVID-19 as describe in **Figure 1**, either antisense would bind to genomic RNA templates or discontinuous transcripts of virus which would inhibit the viral protein translation and virions reproduction ultimately. In order to regulate target gene expression, antisense ought to reach disease-associated tissues and cross cell membranes. This is in part facilitated by the manipulation of their chemical structure, which makes oligonucleotides also stronger and safer with a lower chance to have side effects on host immune system(1–3). Apart from this, the most important factor for RNA virus is that we need highly stable and frequent region where your antisense would bind(5–7). Based on this idea, we decided to count frequency of each plausible target via most frequent region(MFR).

**Figure 1 :**
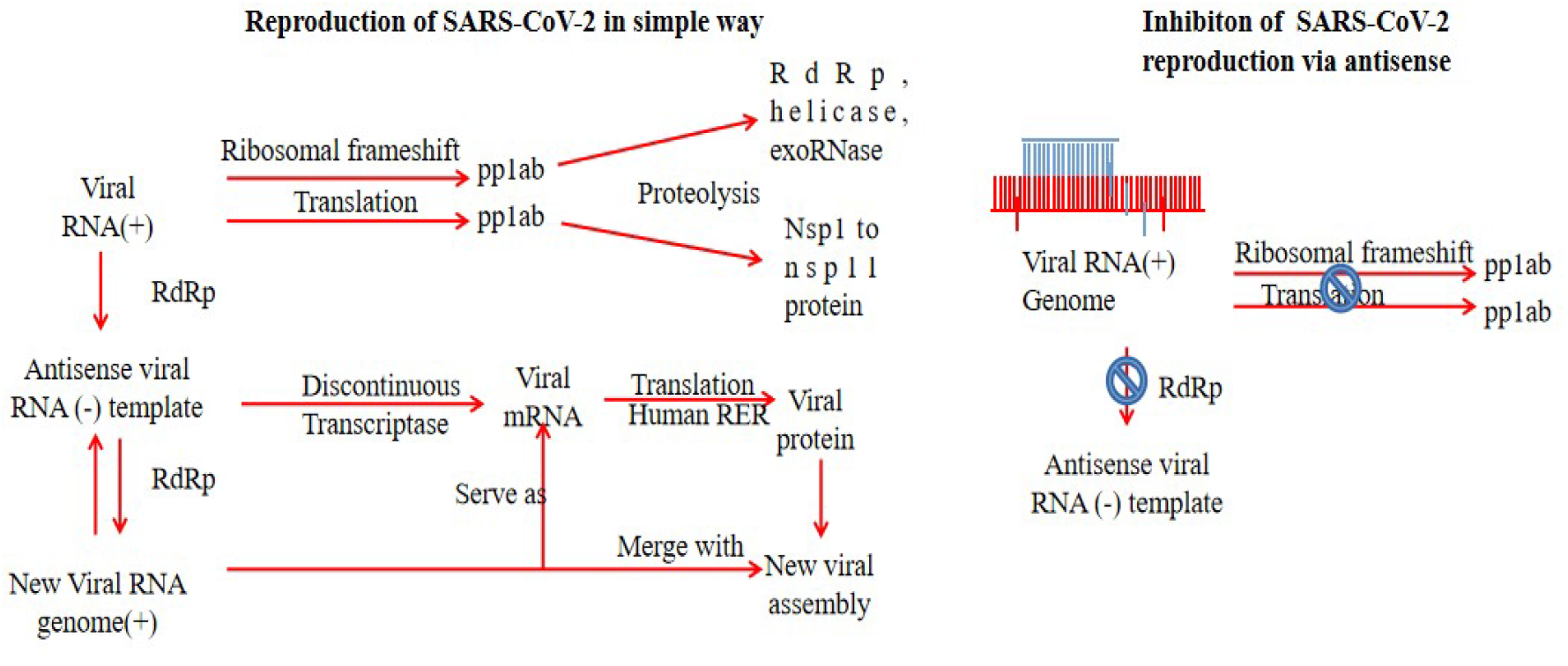
The schematic diagram is showing reproduction of SARS-CoV-2 in simple way(Left side) and how antisense could be helpful to inhibit the SARS-CoV-2 reproduction via antisense blocking. Antisense binds complementarily to inhibit translation and negative template synthesis via RdRp protein.

Based on literature survey, the number of algorithm or method were available to find conserved region or homology. For example, Clustal Omega multiple sequence alignment tool(8,9) and NCBI BLAST(10). This tool mostly use for finding conserved regions. But this tool has certain limitation i.e. it can process only 4000 sequences or 4MB size file(8,9). The number of open source software around us to design antisense and siRNA for all such as OptiRNAi(11), siExplorer(12), DSIR(13), i-Score(7), MysiRNA-Designer(14), siDirect(15), siRNA Selection Server(16), siRNA-Finder(17), PFRED(18) and OligoWalk(19). These are all web based design tools. These algorithms predict and aid in designing/development of best antisense but not considers frequency as major finding criteria. And therefore we designed DURVA and MFR methods. And we initially hypothesized that this new approach would be helpful to find out highly conserved region from SARS-CoV-2 RNA sequences. And based on hypothesis and objectives, we designed scripts of DURVA and MFR method using Python programming language. Our first objective is to find regions which are meeting Chan et. al(3) criteria for antisense designing. Based on our study, we understood that DURVA - MFR can process more than ten thousand of 30kbp long sequences, and can calculate most frequent region from 1Gb text file size. At the end of our study, we concluded that our defined ASO-target sites are more than 88% identical with SARS-Cov-2 data. This could be helpful to understand how our method will be useful in future to find most frequents and highly conserved target site from large sample size.

## Methods and Tools

Based on our short piolet study for finding most frequent region, we modified our python transcripts to process large sample size in short turnaround time. These transcripts are very easy and can work on Linux and Windows operating systems. To design and run the python scripts we used IDLE Python3.9 version on Microsoft windows11. The DURVA MFR method work as followed:

### 1) Simplifying whole genome sequence in single line

To find most frequent target, we aggregated all our previous data. To check the efficiency of python script to convert whole genome in single line format for sequences downloaded from NCBI Virus database (https://www.ncbi.nlm.nih.gov/). Further, we had downloaded 13875 SARS-CoV-2 RNA sequences from NCBI Virus database(20) (https://www.ncbi.nlm.nih.gov/). In which 4144 were Asian sequences, 2024 were European sequences, 2266 were African sequences, 2427 were Kiwis (New Zealand) samples, 1185 were North American samples and 1829 were South American samples. We removed newline character (\n) from all this samples for making DURVA method simple. Newline character (\n) works same as ‘Enter’ in our text file.

### 2) Deletion of unwanted region from virus sequence alignment (DURVA)

In this step, we studied 13875 SARS-CoV-2 RNA sequences (https://www.ncbi.nlm.nih.gov/) (4144 were African sequences, 2427 were Kiwis (New Zealand) sequences, 1185 were North American sequences and 1829 were South American sequences). We deleted unwanted region from all these sequences according to Chan et. al(3) antisense guidelines.

### 3) Most frequent antisense target region (MFR)

In piolet study, we went through processing high number of sample in short interval. So, we improvised MFR python script to find most frequent region irrespective of comparing known reference antisense target. Herein, we modified Severance logic for frequency. After improvisation, we included both complete and partial genome sequences for the study. we calculated frequency of targets from all these 13875 SARS-CoV-2 RNA sequences to find most frequent target regions.

### 4) Designing and development of antisense sequence

As per Watson-crick DNA structural model, the antisense would bind complementary to mRNA. And designed antisense would bind to target sites which we found as outcome of MFR method. The short and very frequent sequence would be helpful to design and development of antisense. Thus; the antisense would have complementary nucleotides according to Watson-Crick DNA model. And on the principle of this assumption, we wrote our python scripts for developing antisense sequence from 5’⟶3’ direction.

Initially, we designed antisense sequences based on all MFR data. Continental and world-wide level. And we found the best antisense from previous SARS-CoV-2 complete and partial genome sequences for piolet study. Finally, we modified our result by selecting loci of target in genome of SARS-CoV-2. After improvisation, we included both complete and partial genome sequences for the study.

## Data analysis

The frequency of each target calculated via MFR method and based on MFR outcomes, we designed antisense by applying reverse complementary python code for each target. Similarity or homology between multiple sequences and targets were calculated via Jalview software (Clustal Omega) (8,9) and NCBI BLAST (https://blast.ncbi.nlm.nih.gov/Blast.cgi)(10). The statistical data collected for parameters such as frequency, similarity, free energy and temperature etc. Data analysis and graph plots were generated via GraphPad Prism 8.0.1 version. The hybridisation thermodynamic were evaluated via online Oligowalk (siRNA web server)(19) (https://rna.urmc.rochester.edu/cgi-bin/server_exe/oligowalk/oligowalk_form.cgi) and bimolecular secondary structure between antisense and target were predicted via RNA structure webserver(21) (https://rna.urmc.rochester.edu/RNAstructureWeb/Servers/bifold/bifold.html)etc.

## Results and discussion

Designing ASOs and siRNA can be logically challenging for selection of tool or therapeutic sequences. There are number of online open-source software around us to design antisense and siRNA such as OptiRNAi(11), siExplorer(12), DSIR(13), i-Score(7), MysiRNA-Designer(14), siDirect(15), siRNA Selection Server(16), siRNA-Finder(17), PFRED(18) and OligoWalk(19). But in our study, we discussed only those tools which are helpful to find similarity and stability of antisense target or antisense relevant to our proposed method such as NCBI BLAST(10), Jalview (Clustal-Omega Multiple alignment tool) (EBI) (8,9) and OligoWalk(19) etc.**(Table 1)**

**Table1.**
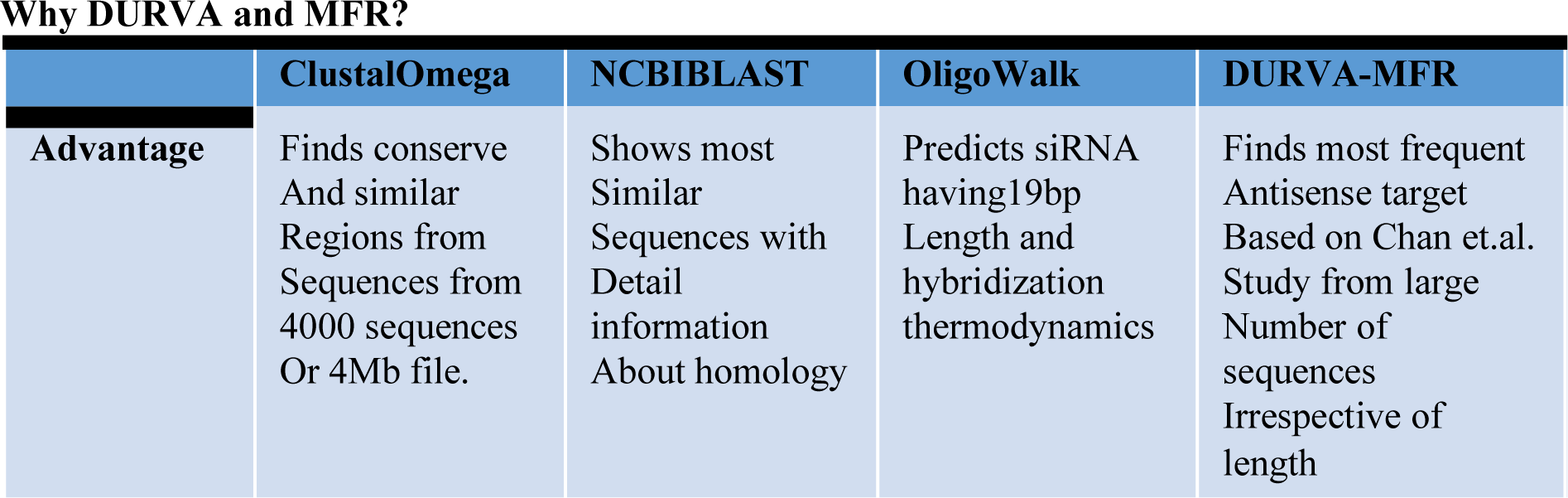
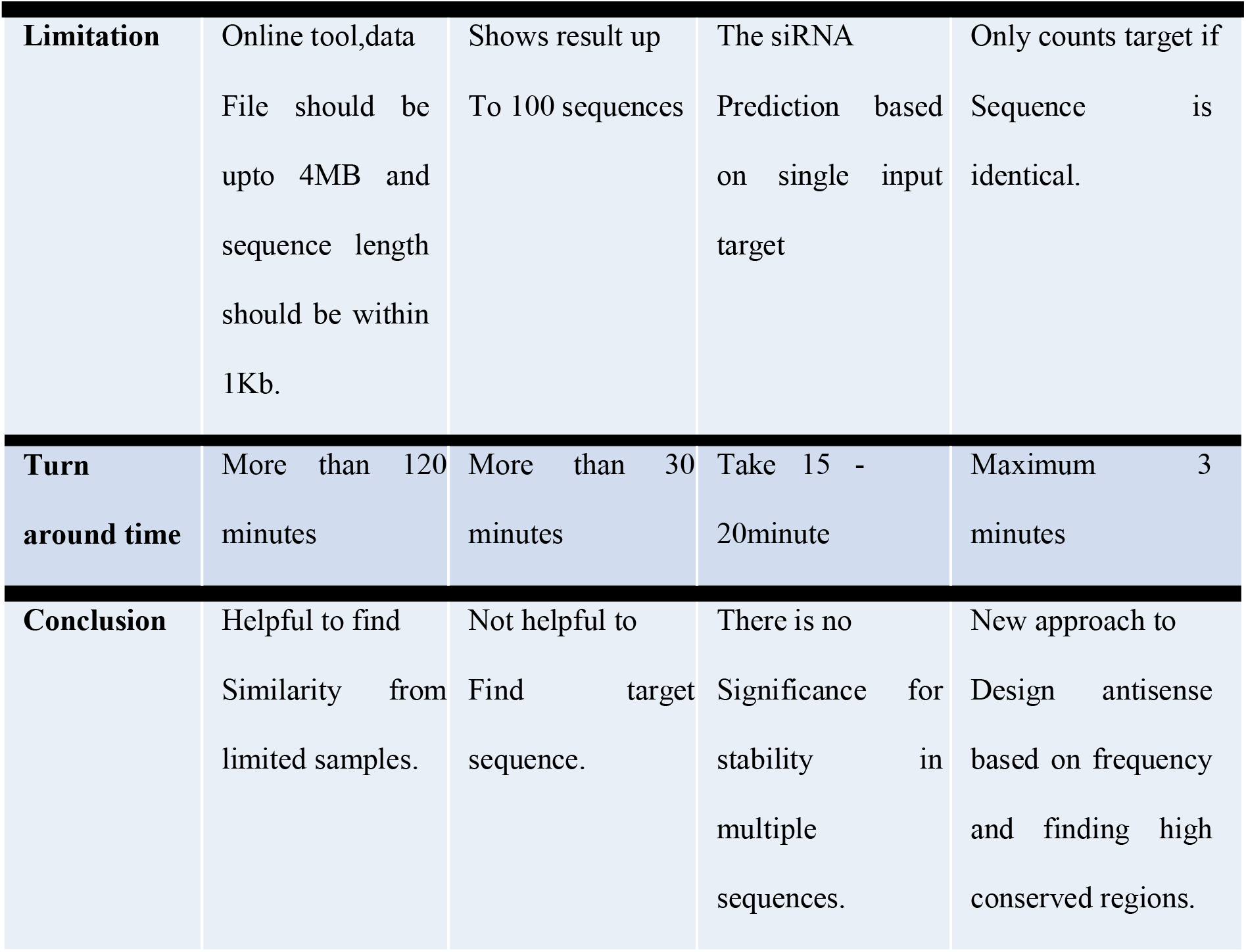
Why DURVA and MFR? Comparative analysis of Clustal omega(8,9), NCBI BLAST(10), OligoWalk(19) and DURVA-MFR. The comparison between tools based on advantage, limitation, turnaround time of tools or method. We also defined the prime use of tools in particular aspect for design and development of antisense.

|  | ClustalOmega | NCBIBLAST | OligoWalk | DURVA-MFR |
| --- | --- | --- | --- | --- |
| **Advantage** | Finds conserve And similar Regions from Sequences from 4000 sequences Or 4Mb file. | Shows most Similar Sequences with Detail information About homology | Predicts siRNA having19bp Length and hybridization thermodynamics | Finds most frequent Antisense target Based on Chan et.al. Study from large Number of sequences Irrespective of length |
| **Limitation** | Online tool,data File should be upto 4MB and sequence length should be within 1Kb. | Shows result up To 100 sequences | The siRNA Prediction based on single input target | Only counts target if Sequence is identical. |
| **Turn around time** | More than 120 minutes | More than 30 minutes | Take 15 - 20minute | Maximum 3 minutes |
| **Conclusion** | Helpful to find Similarity from limited samples. | Not helpful to Find target sequence. | There is no Significance for stability in multiple sequences. | New approach to Design antisense based on frequency and finding high conserved regions. |

In our study we are working on novel and simple simulation technique which is effective and producing data in short time. For processing the samples we used IDLE Python3.9 versions on Microsoft windows11 to write transcripts for DURVA MFR.Workflow of DURVA and MFR as follow as shown in **Figure 2**.

**Figure 2:**
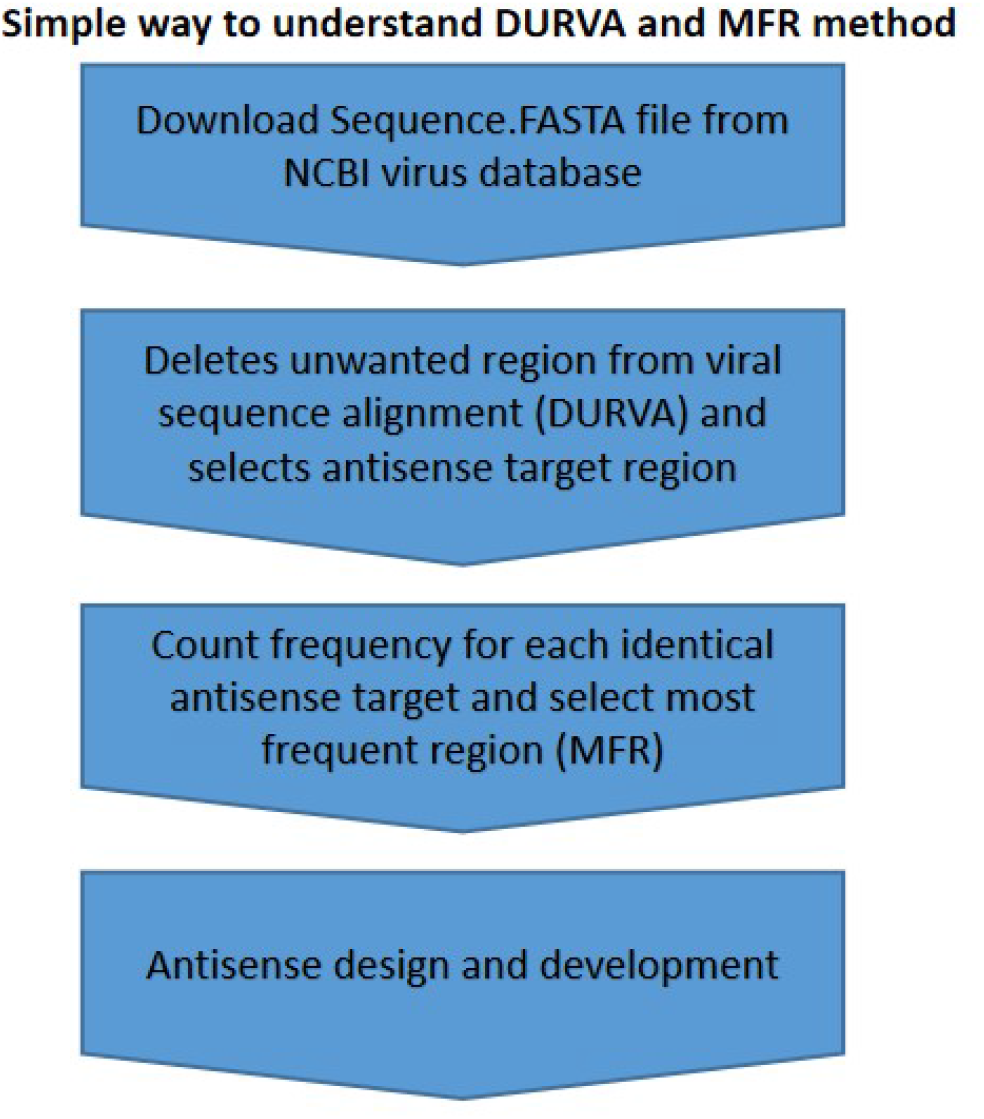
This is simplest block diagram to understant how DURVA and MFR will helpful to simulate targets and antisense sequence design for lethal infection. The detail about process were explained in method and tools section.

### 1) Simplifying whole genome sequence in single line by removing newline character

In piolet study, we checked the efficiency of this transcript to convert whole genome in single line format for high number and long sequences in very few seconds. After removing newline character from each end of 13875 complete or partial sequences made further process easy. Each line without new line character serves as one word which contains equal number of characters or letters same as the length of each sequence.

### 2) Deletion of unwanted region from viral sequence alignment(DURVA)

After removing newline character (\n) from whole genome sequence and simplifying the genome sequence study, it eased our primary aid to delete unwanted region from all virus sequences according to Chan et. al(3)criteria. We studied 13875 SARS-CoV-2 RNA sequences (SARS-Cov-2 targets.txt). We deleted unwanted region from all these sequences to find most frequent target region.The DURVA extracted and stored 100 to 4000 targets from each sample sequence. To evaluate speed of DURVA, we aggregated all continental data in single file and ran the program file. We concluded that the DURVA shortlisted and stored desirable target sequences within few minute for high number of sample sequences. This proved the efficiency, accuracy and shorten the time length for DURVA method.

### 3) Most frequent antisense target region (MFR)

This was most important and challenging step in our critical thinking which could narrate our all results and accuracy to find most frequent target region. In preliminary study, we matched our selected sequences with respective to reference sequence (NC_045512.2) https://www.ncbi.nlm.nih.gov/)(10,20). The previous thinking was lengthy and not working properly and harmful for desktop functioning. Although, some of the targets were not identical with targets of reference sequences after MFR run. So; MFR did not count those targets or samples indirectly.

Recently, we improvised MFR python script to find most frequent region by using Severance logic. This improved method found most frequent antisense target from DURVA data **(Table2)**. In this improvised process we need not to compare unknown target sequence with known reference sequences (NC_045512.2) https://www.ncbi.nlm.nih.gov/)(10,20). antisense target sequence. This would not only find certain most frequent targets, also helpful in finding highly stable region throughout the studied sequences irrespective of loci. However; the advantage of this improvisation was: it reduced turnaround time and increased our accuracy. Also; the dependency of other tools such as Oligowalk and NCBI virus became lesser than previous strategy. After improvisation, we included both complete and partial genome sequences and compared old data with new results to evaluate our hypothesis. And it showed that the decrease in frequency percentage as we seen in South American population in pilot study. The reason behind that the decrease in frequency because a)some of the target region were belonged to other gene or b)contain unnecessary characters instead of nucleotides i.e. NNNNNN. However, the small change in previous script lead us to remarkable results as shown **(Table2)**.

**Table2.**
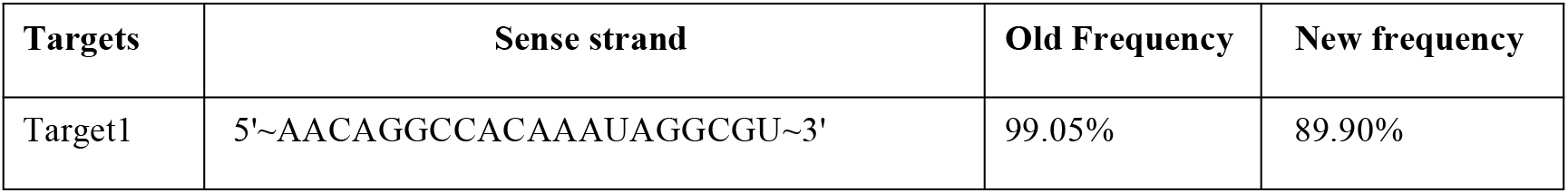
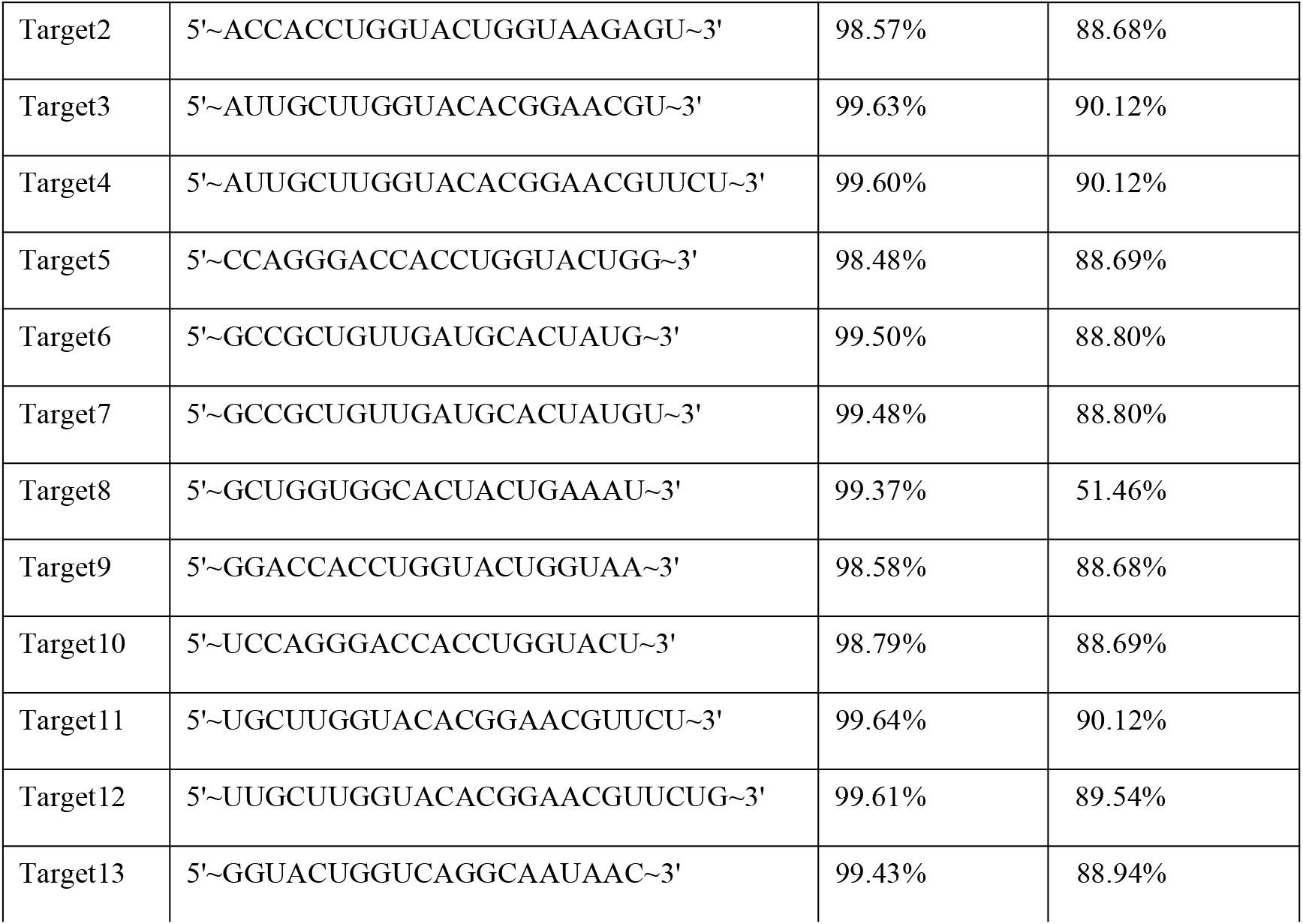
Comparison of modified MFR python scripts between new and older frequencies. The data represents most frequent targets discovered from SARS-CoV-2 genomic RNA sequences. These thirteen targets discovered from a)n=19572SARS-CoV-2 sequences for old frequency and b)n=13875SARS-CoV-2 sequences for new frequency.

### 4) Designing and development of antisense sequence

As per Watson-crick DNA model, the antisense would bind complementary to mRNA. And developed antisense would bind to target site which we discovered as outcome of MFR method. The short and very frequent sequence would be helpful to design and development of antisense that were acquired to fight against SARS-CoV-2 virus.

In short, DURVA and MFR is novel and simple approach for designing antisense because Jalview (Clustal-Omega Multiple alignment tool) (EBI) (8,9) did not deal with very large numbers (tens of thousands) of DNA/RNA or protein sequences due to it use of them BED algorithm for calculating guide trees. DURVA-MFR can be alternative and very simple method to find most identical and probable conserve region within maximum number of sequences by ignoring transcriptional errors. Similar to Clustal omega Multiple sequence alignment, T-coffee and Lalign(8,9) are used to find local alignment not to design target as per antisense requirement. Also; these tools can process if your input file contains 500 sequences or having size of1Mb. While DURVA and MFR can study more than ten of thousand sequences and up to 1Gb file. This is more accurate and less challenging to design therapeutic ASO without adding available features related to homology. We are already familiar with NCBI BLAST(10) and Oligowalk(19) web server tools for homology and antisense prediction. But, we also discussed limitation and turn around time of these tools in **Table3**. While the advantage of DURVA-MFR method over the finding efficient and high conserved region without need of BLAST(10) for homology and OligoWalk(19) for prediction of antisense. Because the target discovered from high number of sample size or annoted sequences of same species which were available at NCBI virus database (https://www.ncbi.nlm.nih.gov/)(10,20). And the Oligowalk SiRNA web server(19) predicts the efficient siRNA based on single target input with limit while DURVA-MFR method can define most efficient antisense based on frequency of target analysed from large number of long sequences.

**Table3.**
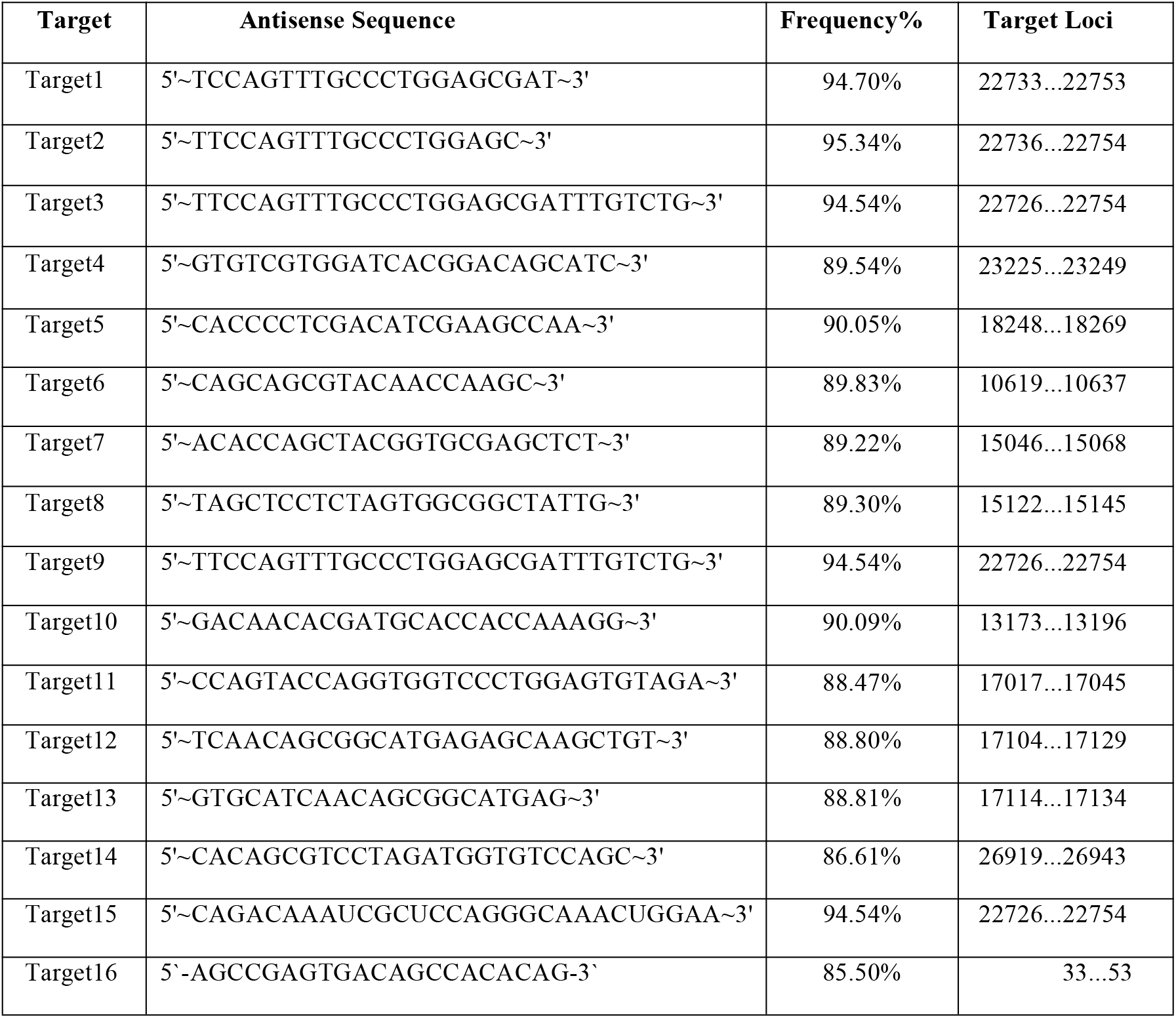
Antisense sequence designed based on modified method. These are top 16 designed antisense sequences (including Target16 is previous potential antisense) with frequency and target loci on genomic RNA of SARS-CoV-2.

The regression analysis with few studies related to potential ASO designed via high-score from oligonucleotide properties calculator tool. Our analysis proved the success rate of our newly innovated method. Goryachev and co-workers did comparative analysis of old and new SARS strains. Based on analysis they tested synthetic ASO i.e. 5’- AGCCGAGTGACAGCCACACAG-3’ within in vitro model(1). The stability of their antisense template with respective to our findings was very lesser as comparison. The previous discovered sense strand was 85.5% frequent within our SARS-CoV-2 genomic RNA sequences data **(Graph 3)**. The previous verified ASO target site was less than 86% frequent with SARS-Cov-2 data while our defined all ASO target sites are more than 86.61% frequent with SARS-Cov-2 data. However, both studies indicate that ORF1ab is most stable gene within new strains of corona virus. This completely states that combination of DURVA and MFR not only efficient to find most frequent region but also helpful for designing potential ASO. This analysis would be helpful for upcoming clinical trials against this worldwide pandemic. Moreover, H. Chubuk study for promising strategy against SARS-CoV-2 defined AON against 5’UTR, 3’UTR and start codon based on high-score of parameter described by Aartsma-Rus et al. He also proposed that targeting viral genome could be helpful to control viral replication and infection(23). However,we compared the frequency of high-scored ASO via DURVA-MFR method. We concluded that frequency of high-scored ASO were between 47-87%. Recently, Yan et al developed structure based ASO to inhibit SARS-CoV-2 replication(2). Their approach would be helpful to cross-check the binding of our designed antisense in 3D space at molecular level.And also helpful to find most useful ASO based drugs against disease like SARS-CoV-2. However, their study was focused on structure in stead of stability of genomic regions. And structure could be varying at molecular level which could be limitation of their critical thinking. However, we predicted bimolecular secondary structure via RNA structure web server (https://rna.urmc.rochester.edu/cgi-bin/server_exe/oligowalk/oligowalk_form.cgi)(21). And we noticed promising RNA strucure bifold results for DURVA and MFR outcomes. The highest value of energy for previously studied high-scored AON and TR_3 was -31.5(23) while lowest value of energy for our frequent antisense and target15 was-37.2.This could be helpful to understand binding efficiency of our discovered antisense.

In our preliminary study, we faced difficulties to find more appropriate outcomes. So; later we improvised MFR python script to find most frequent region irrespective of comparing known reference antisense target by modifying Severance frequency strategy(22). This improved method found most frequent antisense target from DURVA data. In this modified version, we need not to compare unknown target sequence with known reference sequence (NC_045512.2) (https://www.ncbi.nlm.nih.gov/)(10,20) antisense target sequence. This not only found certain most frequent target also helpful in finding highly stable region through out the studied sequences irrespective of loci. In Asian samples, we found that more than 99.28% frequent in all outcomes and the one short target sequence was having high frequency about 99.87%. Contrary Indian study showed decrease in frequency percentage as we seen earlier. There we found only few target sequence with more than 95% frequency. The reason behind the decrease in frequency was either some of the target region were belonged to the gene or having the inappropriate nucleotides (e.g. NNNNNN) at same loci or position. After improvisation, we included both complete and partial genome sequences for the study. However; the benefit of this improvisation was that it does not saved our time only but increased our accuracy too and can process up to 1Gb data. Moreover, the dependency of other tools became lesser comparative to previous strategy. Although, it did not alter all previous results.

These are designed and developed antisense sequences and their target **(Table 3)** was verified by homology were evaluated using NCBI BLAST(10) (https://blast.ncbi.nlm.nih.gov/Blast.cgi), similarity via Jalview Clustal omega multiple alignment tool(8,9) (https://www.ebi.ac.uk/Tools/msa/clustalo/) and stability and hybridisation thermodynamics of most frequent targets OligoWalk tools(18,21) (https://rna.urmc.rochester.edu/cgi-bin/server_exe/oligowalk/oligowalk_form.cgi) respectively.

### DURVA-MFR and other tools

However, the major drawback of this tool is: if any error in sequences due to improper sequencing or mutation, it does not able to find conserve regions from more sequences or complete genome of SARS-CoV-2 virus in such cases. As some of the sequences did not contain 100% identical fragment or complete fragment with respective to reference fragment after MFR run. So; MFR did not counts those samples. Therefore,we chose Jalview (Clustal-Omega Multiple alignment tool) (EBI) (8,9) and NCBI BLAST(10).These tools helped us to find similarity between partial or semi-conserved fragments and most frequent region within non-identical antisense-target sequences. This helped to assimilate relation between DURVA-MFR and Clustal omega. Clustal-Omega Multiple alignment tool (8,9) results showed that most frequent regions were highly similar with few genomic RNA sequences and BLAST(10) results had shown all short template having E value<0.006 against SARS-CoV-2. The **Supplementary file(Target.xls)** contains data for partially or semi-conserved with these 16 targets. The statistical data analysis was done in column and grouped table type and nested graph type for data analysis to generate graph1, graph2 respectively via GraphPad Prism 8.0.1 version. The graph 1 and graph 2 defined the similarity in terms of PID (8,9) and comparative analysis between identical and similarity in terms of percentage respectively. As per our observations in terms of PID, Target1, Target3, Target4, Target9 and Target15 were identical within most frequent genomic RNA sequences and most similar within few RNA sequences too.

And here we compare oligoWalk(19) data for parameters such as stability and efficacy of our discovered antisense against previously known anstisense. In short, hybridisation thermodynamic and probabilities of efficient siRNA. Here we compared our discovered antisense with similar siRNA predicted via oligoWalk(19). The similar siRNAs are having efficient siRNA probabilities more than 0.55. Six antisense complementary against Target4, Target6, Target7, Target10 and Target11 were having efficient siRNA probabilities more than 0.8. Similar siRNA sequence had overall free energy (ΔGoverall) in the range of -17.4 kcal/mol to -28.1 kcal/mol and mean -22.04 kcal/mol. The free energy required for duplex formation (ΔG_Duplex_) in the range of -32.7 kcal/mol to -37.7 kcal/mol and mean -34.25 kcal/mol. The temperature (Tm-Dup) required for duplex formation in the range of 82.4^0^Cto 91.1^0^C and mean 85.74^0^C. The free energy to break-target (ΔGtarget) in the range of -4.7 kcal/mol to -14.7 kcal/mol and mean -10.19 kcal/mol. The free energy of intra oligo (ΔG_intra-oligomer_) in the range of -0.1 kcal/mol to -3.3 kcal/mol and mean -1.00 kcal/mol and inter oligo(ΔG_inter-oligomer_) in the range of -10.6 kcal/mol to -15 kcal/mol and mean -12.04 kcal/mol. And free energy between 5’ and 3’ ends of predicted functional siRNA (ΔGEnd_diff) in the range of -0.73 kcal/mol to 2.32 kcal/mol and mean -1.44 kcal/mol.

Nevertheless, we compared our result with one of potential therapeutic ASO tested against new SARS strains within in vitro experiments. The synthetic ASO i.e. 5’- AGCCGAGTGACAGCCACACAG-3’ and sense strand 5’- CTGTGTGGCTGTCACTCGGCT-3’ with respective to our studied parameters. After our conclusion we could state that their discovered sense strand was 85.5% frequent within our SARS-CoV-2 genomic RNA sequences data. The more common thing we have noticed within both experiments after the BLAST of most frequent region i.e. 5’∼UGCUUGGUACACGGAACGUUCU∼3’ that it also belongs to ORF1ab gene of SARS-CoV-2 as same like 5’-CTGTGTGGCTGTCACTCGGCT-3’. This both studies suggest that ORF1ab is most stable gene within new strains of corona virus based on analysis. Also; we have calculated hybridisation thermodynamic and probabilities of efficient siRNA for known potential antisense(1). The similar siRNA 5’-AGCAUGCAGCCGAGUGACA-3’ sequence with respective to known potential antisense sequence 5’- AGCCGAGUGACAGCCACACAG-3’ were having efficient siRNA probabilities 0.33, ΔGoverall were 22.5 kcal/mol, ΔGDuplex were -37.9 kcal/mol, Tm-Dup90.8^0^C, ΔGtarget were-11.2 kcal/mol, ΔGintra-oligomer were -1.4 kcal/mol, ΔGinter-oligomer were -16.6 kcal/mol and ΔGEnd_diff was 0.03kcal/mol etc.

By comparing oligowalk data(19) for paramters such as melting temperature, free energies such as overall, interoligo and intraoligo and break target etc. for previous against antisense designed via DURVA and MFR suggested that our discovered antisense were having more efficient siRNA probabilities compared to known potential antisense. Six similar antisense templates (against Target4, Target6, Target8, Target10, Target13 and Target14) were having more negative ΔGoverall. It meant tighter binding against genomic RNA and transcribed RNA template of virus. One similar antisense templates (against Target8) was having ΔGDuplex and T_m-Dup_ values near to known potential antisense. It meant both were equally more stable and requires high melting temperature to hybridise duplex formation. Eight similar antisense template (against Target1, Target2, Target3, Target5, Target7, Target9, Target11 and Target12) were having more negative ΔGtarget. It meant these antisense templates will be less accessible for siRNA binding. Four similar antisense template (against Target8, Target11, Target13 and Target14) were having more negative ΔGintra-oligomer values. It meant these are more self-stable structure comparative to known potential antisense template. All similar discovered template via DURVA-MFR were having more positive ΔGEnd_diff. This meant that all similar functional siRNA will be shown an unstable 5’end. Contrary, 5’-AGCAUGCAGCCGAGUGACA-3’ was more stable dimerization comparative to our discovered antisense templates.

The multiple variable table type and column graph type plotted via GraphPad Prism 8.0.1 version for data analysis of graph3. Graph3 is correlation between parameters such as frequency, similarity, melting temperature required for duplex formation and free energies such as ΔGoverall, ΔGDuplex, ΔGtarget, ΔGintra-oligomer, ΔGinter-oligomer and ΔGEnd_diff. Overall data suggested that DURVA and MFR method of finding antisense work efficiently and accurately and can be helpful for designing and development of antisense.

Also; we studied predicted bimolecular secondary structure derived from RNA structure web server (https://rna.urmc.rochester.edu/cgi-bin/server_exe/oligowalk/oligowalk_form.cgi)(21). And we noticed all promising RNAstrucure bifold results for DURVA and MFR outcomes. The highest value of energy for previously studied high-scored AON and TR_3 was -31.5(23) while lowest value of energy for our frequent antisense and target15 was -37.2. This could be helpful to understand binding efficiency of our discovered antisense. But except target15, remaining targets and antisense were binding 100%. These predicted bimolecular secondary structures can be helpful to understand how our discovered antisense will best for controlling SARS-CoV-2 replication.

### Future scope of DURVA and MFR method

Apart from antisense designing and developing DURVA-MFR method can be used in finding most stable homologues region between two or more sequences. Using these strategies, we could find similarity between paralogue and orthologue sequences. These algorithms have ability to cut short sequences in serial. This serial sequences can be compared with remaining other sequence and can evaluate most identical frequent region within two different species or similar species. This method helpful in finding most stable and identical region within mutating virus such as SARS-CoV-2, HIV and influenza viruses. Using these two algorithms we can design primers and probes for various biotechnological and genetic engineering aids. This method will not only efficient in genomic but useful for proteomics too. This method will mostly use for vaccinology in case of mutating viral strains such as HIV and Corona viruses etc. In future, these methods will be game changer and most powerful logic to design most efficient, accurate, user friendly, time saving and safe antisense to cure lethal infections.

## Conclusion

In this study, we studied all types of parameter as per our hypothetical consideration and proved that our self-designed simulation technique i.e. DURVA and MFR methods can find highly identical and stable antisense targets and design antisense template from large number of sequences and large file size up to 1Gb. These methods are novel and simple way for designing and development of the antisense against all types of lethal infections.

## Acknowledgement

The initial design of algorithm was published in Indian Patent Office Journal No. 08/2021 with Application number 202121005964A.

## Conflict of Interest

There is no conflict of interest reported for this case study.

## Notes

### Competing Interest Statement

The authors have declared no competing interest.

